# Turncoat antibodies unmasked in a model of autoimmune demyelination: from biology to therapy

**DOI:** 10.1101/2024.12.03.623846

**Authors:** Reza Taghipour-Mirakmahaleh, Françoise Morin, Yu Zhang, Louis Bourhoven, Louis-Charles Béland, Qun Zhou, Julie Jaworski, Anna Park, Juan Manuel Dominguez, Jacques Corbeil, Eoin P. Flanagan, Romain Marignier, Catherine Larochelle, Steven Kerfoot, Luc Vallières

## Abstract

Autoantibodies contribute to many autoimmune diseases, yet there is no approved therapy to neutralize them selectively. A popular mouse model, experimental autoimmune encephalomyelitis (EAE), could serve to develop such a therapy, provided we can better understand the nature and importance of the autoantibodies involved. Here we report the discovery of autoantibody-secreting extrafollicular plasmablasts in EAE induced with specific myelin oligodendrocyte glycoprotein (MOG) antigens. Single-cell RNA sequencing reveals that these cells produce non-affinity-matured IgG antibodies. These include pathogenic antibodies competing for shared binding space on MOG’s extracellular domain. Interestingly, the synthetic anti-MOG antibody 8-18C5 can prevent the binding of pathogenic antibodies from either EAE mice or people with MOG antibody disease (MOGAD). Moreover, an 8-18C5 variant carrying the NNAS mutation, which inactivates its effector functions, can reduce EAE severity and promote functional recovery. In brief, this study provides not only a comprehensive characterization of the humoral response in EAE models, but also a proof of concept for a novel therapy to antagonize pathogenic anti-MOG antibodies.

## Introduction

B cells can play a dual role in autoimmune diseases. Firstly, they can capture protein antigens with their immunoglobulins in a membrane-bound form called the B cell receptor^1^. These antigens can then be processed and presented to autoreactive T cells, key players in autoimmunity, resulting in their activation. Secondly, activated B cells can differentiate into short-lived plasmablasts or long-lived plasma cells, which secrete the same immunoglobulins in a soluble form, lacking the transmembrane domain, called antibodies^2,3^. During this process, the immunoglobulin genes can undergo class-switch recombination, generating antibody isotypes with different properties^4,5^, as well as somatic hypermutation, creating antibodies with greater affinity for the cognate antigen^6^. Ultimately and accidentally, these antibodies can cross-react with normal body components, leading to their destruction by the complement system and/or Fc gamma receptor (FcγR)-mediated cellular mechanisms^7,8^.

Typical examples of autoantibodies are those directed against myelin oligodendrocyte glycoprotein (MOG) and aquaporin-4. These autoantibodies serve as diagnostic markers for rare neurological autoimmune diseases, namely MOG antibody diseases (MOGAD) and neuromyelitis optica spectrum disorder (NMOSD)^9–13^. In contrast, no specific antibody has been validated as a marker for the most common neurological autoimmune disease, multiple sclerosis (MS)^14^. However, MS is characterized by the presence, in the cerebrospinal fluid, of a mixture of antibodies, detected by electrophoresis as oligoclonal bands^15^. Unfortunately, there is currently no approved drug to selectively eliminate or counteract these antibodies.

Autoantibodies such as anti-MOG are heterogeneous and not functionally equal^16^. Some recognize accessible epitopes, exposed on the cell surface or in the extracellular environment, making them pathogenic^17^. Others target epitopes that are normally inaccessible due to intracellular localization, protein folding, or proximity to the plasma membrane, preventing them from being directly pathogenic^17^. Pathogenic anti-MOG antibodies generate a positive signal in a cell-based assay used to diagnose MOGAD^11,13,18–22^. Accordingly, anti-MOG antibodies that are negative in this assay are thought to be non-pathogenic. These are nevertheless detectable by ELISA or Western blotting in several conditions such as MS^17,23,24^, MOGAD^25^, NMOSD^26^, and genetic leukoencephalopathies^27^. The function of these “non-pathogenic” antibodies, if any, is unknown.

Much of our knowledge on autoimmune demyelination comes from a mouse model called experimental autoimmune encephalomyelitis (EAE)^28^. EAE can be induced by immunization with myelin antigens such as MOG-derived polypeptides. When a short polypeptide is used (e.g. the immunodominant epitope MOG_35-55_, which is presented to T cells by dendritic cells^29^), EAE develops in a B cell-independent manner^30,31^. In this context, some B cells even exert a beneficial role, as their depletion exacerbates EAE^30–40^. These anti-inflammatory B cells (also called regulatory B cells or Bregs) can attenuate EAE via IL-10 secretion^31,37,41,42^. In contrast, when EAE is induced with a longer polypeptide (e.g. mouse MOG_1-125_, corresponding to the extracellular domain), B cells adopt a detrimental role: they process and present antigenic epitopes^43,44^ and secrete IL-6^43,45^, both leading to encephalitogenic T cell activation^46^. Under these conditions, B cell deficiency attenuates EAE^30,47–51^. Proinflammatory B cells are even essential for EAE induction with either human MOG_1-125_ (hMOG)^47,48,52,53^ or mouse MOG_1-125_ carrying the humanizing mutation S42P abolishing the 35-55 epitope (an antigen called bMOG)^54^. These two antigens are termed B cell-dependent because they do not induce EAE if B cells are depleted.

While there is ample evidence that some EAE models involve proinflammatory B cells as antigen presenters and cytokine secretors, little is known about the involvement of autoantibodies other than the observation that immunization with hMOG induces the production of anti-MOG IgG antibodies with pathogenic potential^53,55^. In the present study, we report the discovery of a population of antibody-secreting B cells that transiently proliferate in lymph nodes of mice immunized with bMOG or hMOG. Our objectives were to: 1) characterize these cells comprehensively by single-cell RNA sequencing and mass cytometry; 2) examine how their antibodies can influence EAE by administrating selected recombinant antibodies or bMOG antiserum; and 3) develop an approach to block pathogenic autoantibodies.

## Results

### Prominent expansion of extrafollicular plasmablasts in B cell-dependent EAE

To determine whether antibody-secreting cells are generated in B cell-dependent EAE, we quantified cells expressing the canonical marker CD138 (syndecan-1) by flow cytometry in various tissues and at different time points (days 0, 8, 12, 24) after immunization with bMOG. These cells were significantly increased in number in the draining inguinal lymph nodes on day 8 as well as in the bone marrow and spinal cord on day 24, but not in the spleen at any time point (**Figure 1A** and **1B**). A similar increase was observed in lymph nodes of mice immunized with hMOG (**Figure 1C**), but not with mouse MOG_35-55_ or adjuvants alone (**Figure 1D**). A replicate of the experiment using Vert-X mice, which express EGFP under the control of the IL-10 promoter^56^, revealed that 62 ± 16 % of bMOG-induced CD138^+^ cells were IL-10 producers (**Figure 1E** and **1F**). A similar percentage was observed in mice immunized with MOG_35-55_ or adjuvants alone (**Figure 1F**), suggesting that IL-10 production by CD138^+^ cells is antigen-independent. However, the total number of CD138^+^ cells expressing IL-10 was much lower under these conditions compared to bMOG (**Figure 1G**). For comparison, only 7 ± 4 % of CD19^+^CD138^−^ B cells expressed IL-10 (**Figure 1F**).

**Figure 1.**
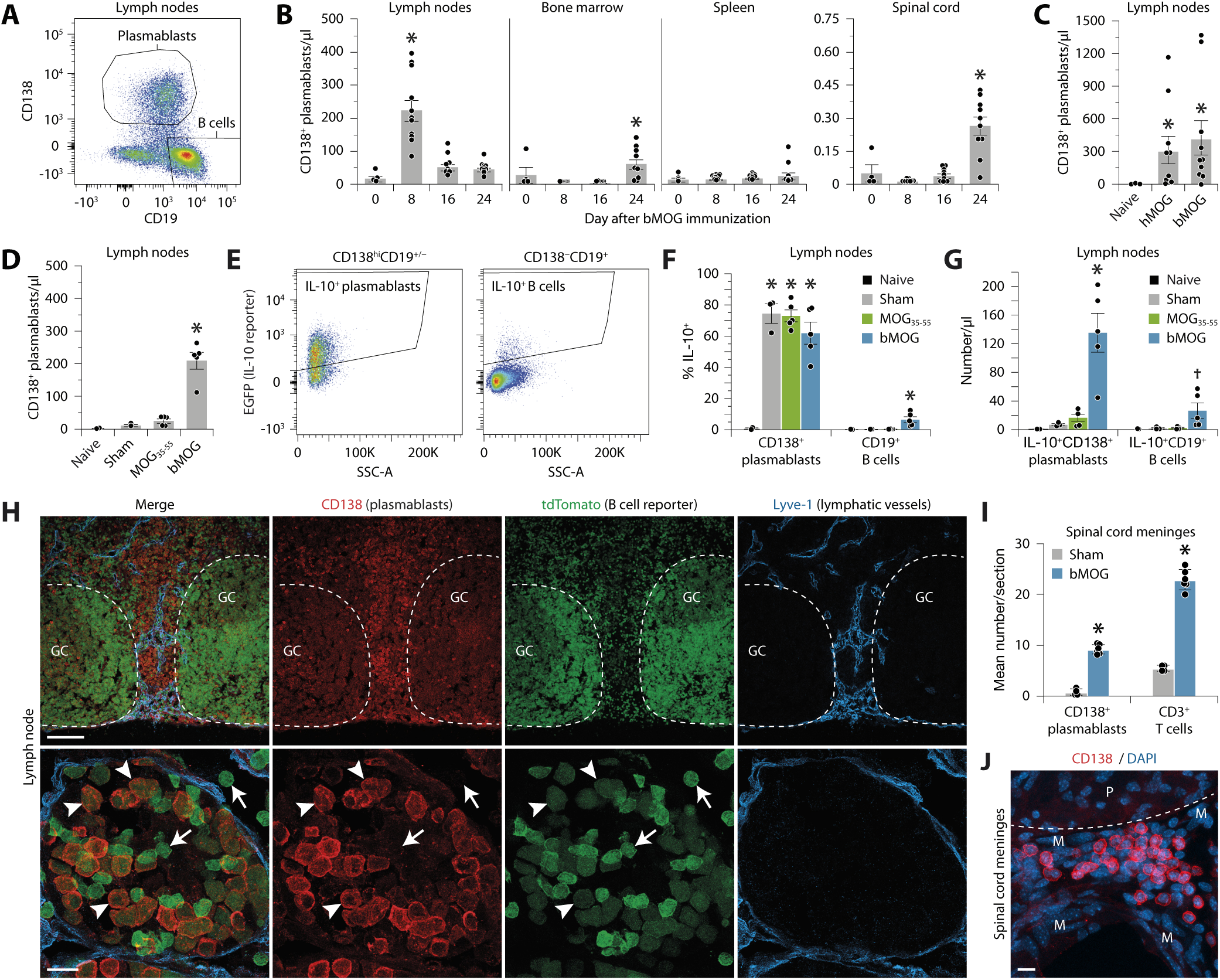
Extrafollicular plasmablasts expand greatly and transiently in EAE induced with B cell-dependent antigens. **(A)** Representative flow cytometry plot of lymph node cells from a mouse immunized with bMOG 8 days earlier. Gating strategy used to identify plasmablasts (CD138^hi^CD19^+/−^) and B cells (CD138^−^CD19^+^) is shown. Dead cells, doublets, and other leukocytes (CD3^+^, Ly6G^+^, CD11b^+^, CD11c^+^) were excluded. **(B)** Quantification of plasmablasts in different tissues and at different time points after bMOG immunization. *Significantly different from the other time points (ANOVA, *P* < 0.0001; post hoc Tukey’s test, *P* < 0.0001). Sample size: 4 day 0, 11 day 8, 10 day 16, 10 day 24. **(C, D)** Quantification of plasmablasts in lymph nodes 8 days after immunization with B cell-dependent (bMOG, hMOG) or -independent (MOG_35-55_) antigens. Mice untreated (naive) or injected with adjuvants alone (sham) were used as controls. *Significantly different from the controls (C: Kruskal-Wallis, *P* = 0.014; post hoc Dunn’s test, *P* ≤ 0.018; D: ANOVA, *P* < 0.0001; post hoc two-tailed Student’s *t*-test, *P* < 0.0001). Sample size: C, 3 naive, 10 hMOG1-125, 10 bMOG; D, 2 naive, 3 sham, 5 MOG_35-55_, 5 bMOG. **(E)** Flow cytometric gating strategy used to identify plasmablasts and B cells expressing the IL-10 reporter EGFP in lymph nodes of Vert-X^+/+^ mice 8 days after bMOG immunization. **(F, G)** Percentage and relative number of plasmablasts and B cells expressing IL-10 in lymph nodes of Vert-X^+/+^ mice after the indicated treatments. *Significantly different from the other groups (ANOVA, *P* ≤ 0.01; post hoc two-tailed Student’s *t*-test, *P* ≤ 0.05). †Tends to be significant (ANOVA, *P* = 0.0588; Student’s *t*-test: *P* ≤ 0.0477). Sample size: 2 naive, 3 sham, 5 MOG_35-55_, 5 bMOG. **(H)** Confocal images at low (top) or high (bottom) magnification showing the distribution of CD138^+^ plasmablasts (red) in a lymph node of a CD19^cre^ × Ai14 mouse expressing the B cell reporter tdTomato (false color green) on day 8 after bMOG immunization. Note that these plasmablasts: 1) are predominantly located near lymphatic vessels (blue) outside the follicles (delineated by dashed lines); and 2) express lower amount of tdTomato than CD138^−^ B cells. Arrowheads: tdTomato^lo^CD138^+^ plasmablasts. Arrows: tdTomato^hi^CD138^−^ B cells. Abbreviation: GC, germinal center. Scale bars: top, 100 µm; bottom, 10 µm. **(I)** Counts of plasmablasts and T cells in the spinal cord meninges from mice sacrificed on day 28 after immunization with bMOG or adjuvants alone (sham). *Significantly different from sham (two-tailed Student’s *t*-test, *P* < 0.0001). Sample size: 4 sham, 6 bMOG. **(J)** Confocal image of a spinal cord section on day 28 post-immunization showing a cluster of CD138^+^ plasmablasts (red) counterstained with DAPI (blue, nuclei) outside the parenchyma (P), within the meninges (M; delineated by a dashed line). Scale bar: 10 µm.

To further determine the anatomical distribution of CD138^+^ cells, we examined confocal images of lymph node sections from CD19^cre^ × Ai14 mice, which express tdTomato specifically in the B cell lineage^57,58^. Numerous tdTomato^+^CD138^+^ cells were observed on day 8 after bMOG injection, mainly outside the germinal centers, in the vicinity of Lyve-1^+^ lymphatic vessels (**Figure 1H**). Furthermore, microscopic examination of spinal cords on day 24 confirmed the presence of CD138^+^ cells, but only in the leptomeninges (**Figure 1I**). Some were individually scattered (not shown), while others were clustered in follicles containing several dozen cells (**Figure 1J**), as observed in people with progressive MS^59–61^ and transgenic mice expressing a MOG-specific B cell receptor^43,62^. Taken together, these results indicate that a population of extrafollicular plasmablasts, phenotypically similar to the IL-10-producing regulatory plasmablasts described in MOG_35-55_-induced EAE^42^, expand to a much greater extent when EAE is induced with B cell-dependent antigens.

### Single-cell transcriptomic and protein profiling of bMOG-induced plasmablasts

To characterize the transcriptome and antibody repertoire of bMOG-induced plasmablasts, we enriched these cells from lymph nodes of four mice on day 8 post-immunization and analyzed them by scRNAseq using 10× Genomics technology. The four gene expression datasets were pooled to produce a single tSNE plot of all the isolated cells that met our quality criteria (**Figure 2A** and **Supplementary Dataset 1**). Using this plot in combination with cell-specific markers, we identified 6 unambiguous subsets of leukocytes (**Figure 2A**). Plasmablasts were distinguished by the expression of *Sdc1* (CD138) and *Tnfrsf17* (BCMA) (**Figure 2B**). A significant proportion of plasmablasts (∼25 %) expressed proliferation markers such as *Birc5* (**Figure 2C**) and several others (e.g. *Mki67*, *Pclaf*, *Ube2c*, *Cdk1*, *Hist1h2ap*, *Top2a*; **Supplementary Dataset 1**). Strikingly, compared with B cells, plasmablasts had upregulated many genes associated with the protein/antibody synthesis machinery, while they had downregulated many genes associated with antigen presentation (**Figure 2D**). This was confirmed at the protein level by mass cytometry, except for MHCII which remained elevated (**Figure 2E−2G**). Furthermore, plasmablasts appear to be poor producers of cytokines, as only *Il15* and *IL10* were detectable, and only in very low amounts (**Supplementary Dataset 1** and **Figure 2D**). Interestingly, they did not express *Aicda* (**Supplementary Dataset 1**), which encodes the enzyme AID, essential for antibody maturation by somatic hypermutation^63^. Consistent with published observations^42,64,65^, they expressed higher levels of *Cd44*, *Cd93*, *Irf4*, and *Prdm1*, while they had lost expression of *Sell* (**Figure 2D**). Overall, our results support the concept that CD138^+^ cells are extrafollicular plasmablasts engaged in a primary antibody response, and not fully matured plasma cells, notably because they still proliferate and express MHCII^2^.

**Figure 2.**
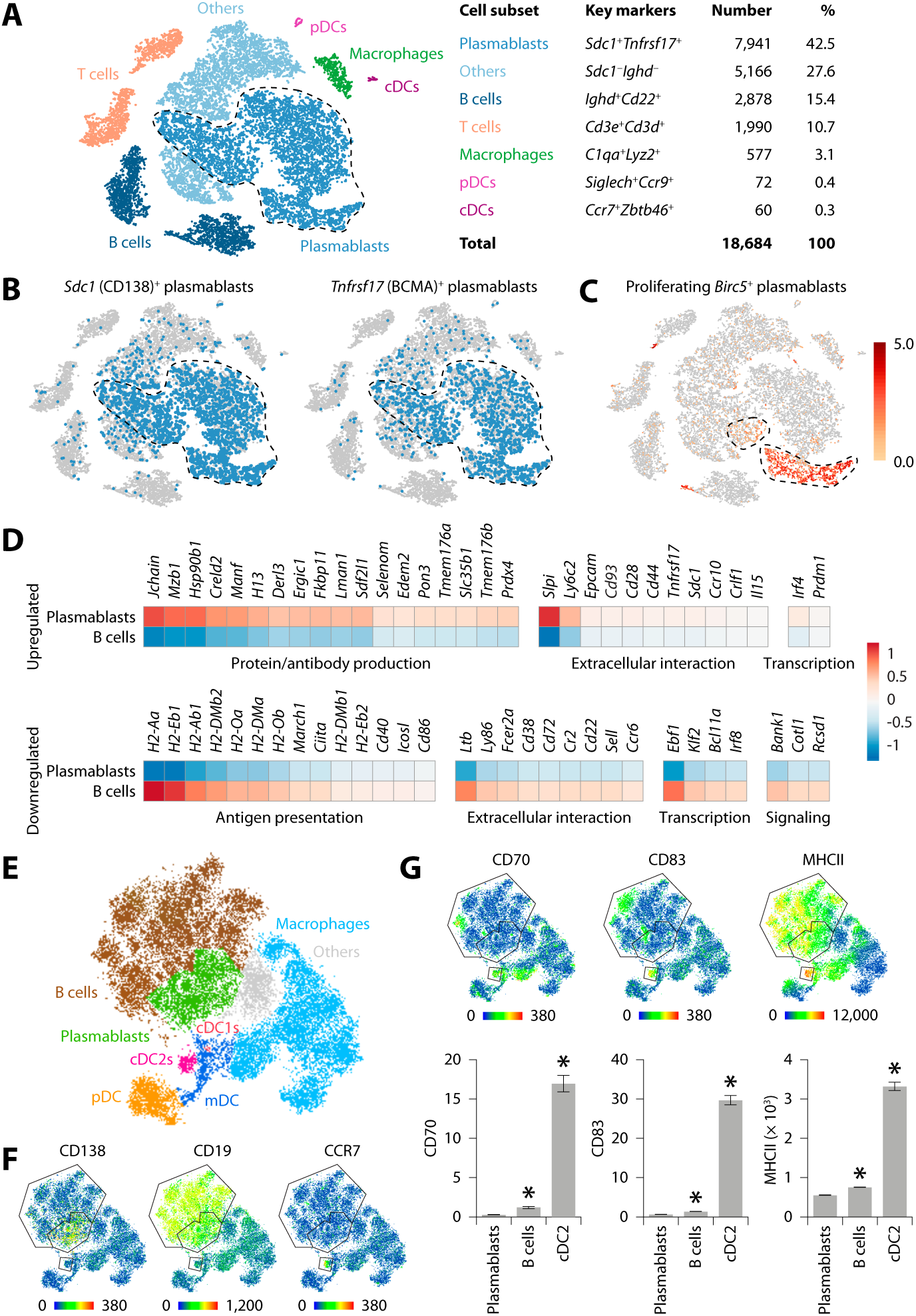
Transcriptomic and protein profiles of bMOG-induced plasmablasts. **(A)** tSNE plot of lymph node cells enriched in CD138^+^ cells from four mice on day 8 after bMOG immunization. Cells were analyzed using 10× Genomics 5’ Single-Cell Gene Expression technology. The indicated cell subsets were identified using the tSNE distribution, K-means, and cell type-specific markers. Encircled cells were those selected for V(D)J profiling in Figure 3. Descriptive data on each subset are provided in table. Abbreviations: pDCs, plasmacytoid dendritic cells; cDCs, conventional dendritic cells. **(B)** Cells expressing the plasmablast markers CD138 and BCMA (blue). **(C)** Log_2_ expression of *Birc5* revealing proliferating plasmablasts (encircled). **(D)** Heat map showing the expression profile of genes differentially regulated in plasmablasts compared to B cells, as identified by significant feature comparison in Loupe Cell Browser (*P* ≤ 0.04). Median-normalized mean UMI counts were Ln(x+1)- transformed and centered without row scaling using Clustvis. **(E)** tSNE plot generated from mass cytometry data on lymph node cells on day 8 after bMOG immunization. **(F)** Markers used to identify plasmablasts (CD138), B cells (CD19), and type-2 cDCs (cDC2s; CCR7) in E. **(G)** Comparison of expression levels of proteins involved in antigen presentation between plasmablasts, B cells, and cDC2s. *Significantly different from plasmablast group (ANOVA, *P* < 0.0001; post hoc Tukey’s test (*P* ≤ 0.0004).

We next analyzed the four V(D)J datasets by selecting only high-quality plasmablasts (**Figure 3A**). Cells with similarly rearranged V(D)J sequences (with or without mutations), most likely arising from a common ancestor, were algorithmically combined into clonotypes, regardless of their constant region (isotypes were considered subclonotypes). Clonotype abundance (**Figure 3B**) and immunoglobulin gene segment usage (**Figure 3C−3E**) were similar between mice. Most clonotypes (93 ± 3 %) had undergone class-switch recombination resulting in the expression of IgG antibodies (**Figure 3D**) lacking the transmembrane exon (**Figure 3F**). IgG1 was the most abundant isotype (75 ± 10 %), followed by IgG2b (10 ± 5 %) and IgG2c (6 ± 3 %). However, these antibodies had not undergone somatic hypermutation, as mutations were infrequent (median = 0; mean = 0.014−0.039 %; **Figure 3G**) and corresponded mainly to junctional additions or deletions in the CDR3 regions, especially that of the heavy chain (**Figure 3H**). Although the clonotype profile was similar between the mice (**Figure 3B−3G**), the percentage of clonotypes shared between at least two mice was only 2.5 % (75 out of 2,974; **Figure 3I**). Among the shared clonotypes, a few had the same V(D)J segments, but different junctional additions or deletions in the CDR3 regions, suggesting that they originated from different ancestors that converged to produce identical or nearly identical antibodies. This was the case for a set of clonotypes, collectively called clonotype 2, which were present in all mice and produced the most prominent antibody, mainly in the form of IgG1 (**Figure 3H** and **3J**). These results indicate that bMOG-induced plasmablasts secrete a complex cocktail of non-affinity-matured IgG antibodies.

**Figure 3.**
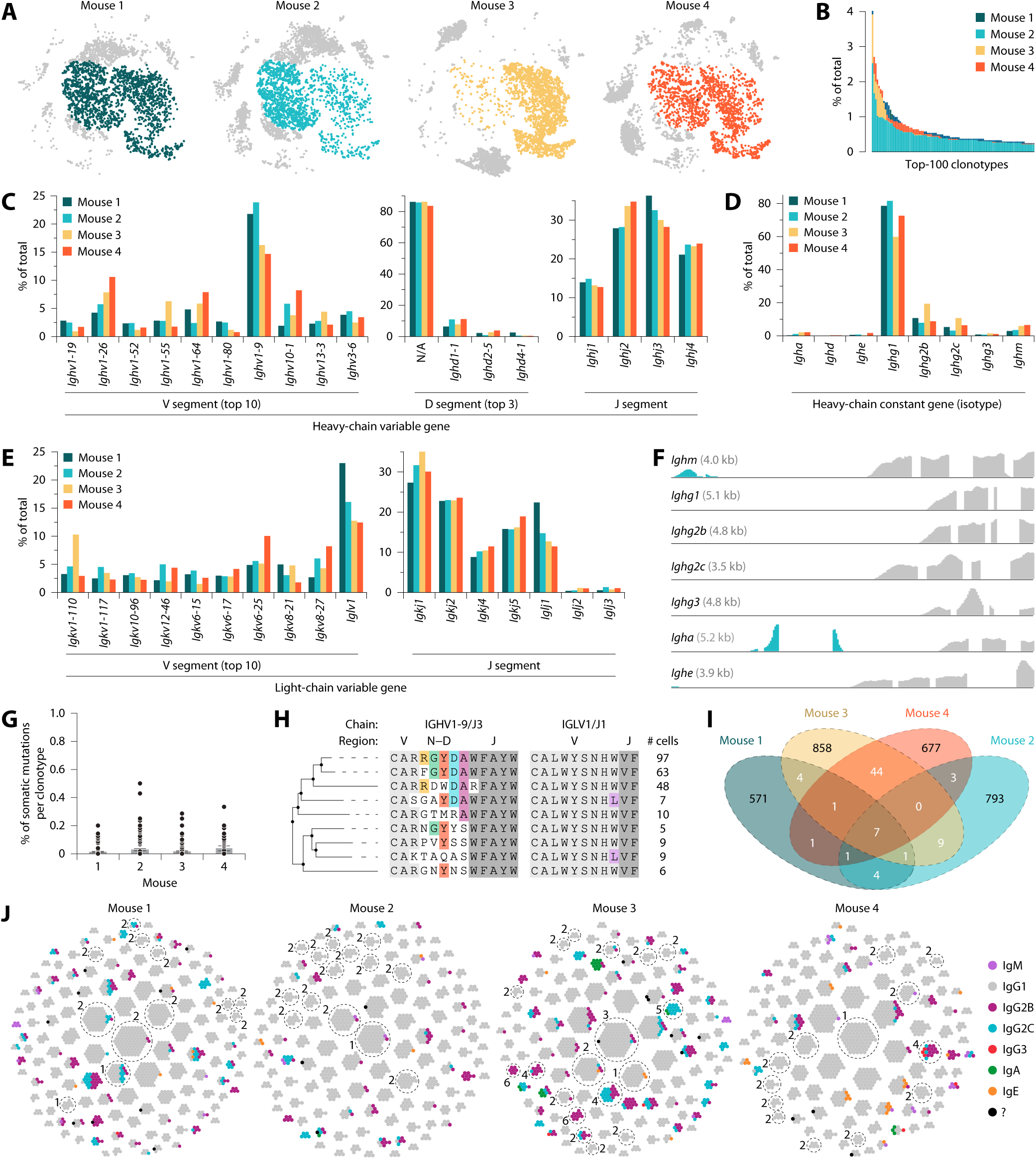
Profile of antibodies produced by bMOG-induced plasmablasts. **(A)** tSNE plots of gene expression data in Figure 2 showing plasmablasts selected for V(D)J analysis (colors). **(B)** Abundance distribution of the top-100 clonotypes, as analyzed using 10× Genomics Single-Cell V(D)J technology. **(C−E)** Comparison of immunoglobulin gene segment usage between mice. **(F)** Sequencing data from gene expression libraries showing the presence of the transmembrane domain (blue) in *Ighm*, *Igha*, and *Ighe*, but not *Ighg* transcripts. **(G)** Percentage of somatic mutations in the *Ighv* gene segment of the main clonotypes with ≥ 5 cells (*n* = 89-123 clonotypes per mouse). Two outliers with mutation rates of 16 and 18 % were excluded in mouse 4. **(H)** Comparison of the CDR3 region of the heavy and light chains of nine highly similar clonotypes from mouse 1, collectively referred to as clonotype 2 in J. These clonotypes were identical except for the CDR3 regions. Alignment was performed with webPRANK. **(I)** Number of clonotypes shared between mice. **(J)** Honeycomb plots showing clonotypes (dot clusters) with ≥ 5 cells (dots) of any isotypes. Numbers indicate clonotypes selected for further analysis.

### Binding capability of recombinant bMOG-induced antibodies

To determine whether bMOG-induced antibodies cross-react with endogenous MOG, we selected 6 clonotypes based on their isotype and size in terms of cell number. Clonotypes 1−3 were the largest and produced mainly IgG1, while clonotypes 4−6 comprised fewer cells, but were the largest to express IgG2b and/or IgG2c (**Figure 3J** and **Supplementary Table 1**). The *Ighv* and *Iglv* gene segments of these clonotypes were cloned to produce the corresponding antibodies (clones C1 to C6) as IgG1, regardless of their original isotype. ELISA results showed that only the IgG1 antibodies (C1−C3) bound to bMOG and that only C1 cross-reacted with mouse and human MOG_1-125_ (**Figure 4A**). Western blots confirmed these findings and showed that C1 could also react with denatured full-length mMOG from spinal cord lysates (**Figure 4B**). However, C1 did not react with either fixed full-length mMOG in spinal cord sections (**Figure 4C**) or native full-length mMOG on live cells in culture (**Figure 4D**), presumably because the epitope is masked due to the protein conformation and/or its proximity to the plasma membrane. Similar results were obtained using C1 in an IgG2b or IgG2c format (**Figure 4D**), ruling out an effect of the isotype on the binding capacity.

**Figure 4.**
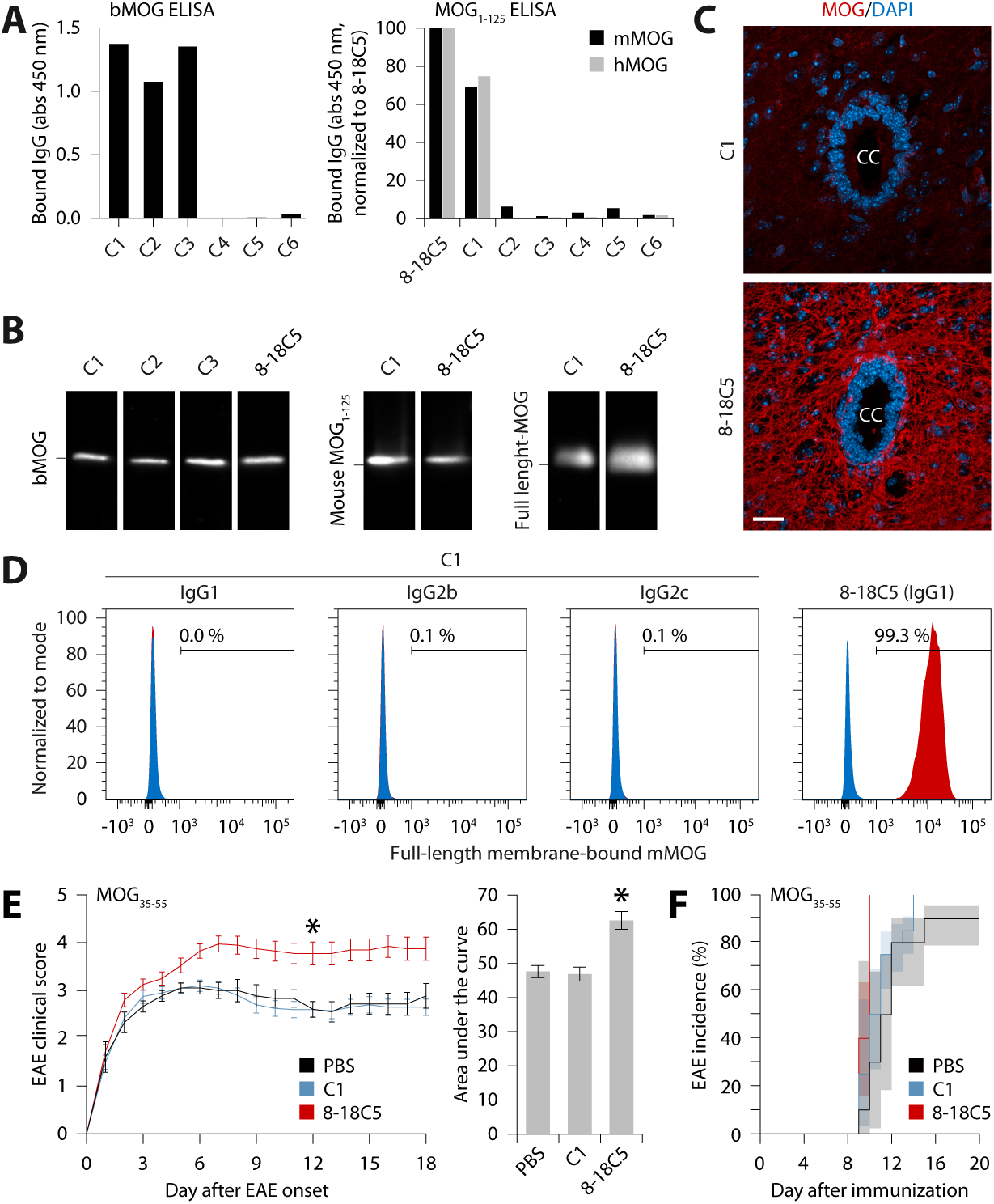
C1 can react with wild-type MOG, but not when the latter is in its native, plasma membrane-bound form, making it a non-pathogenic antibody. In all panels, anti-MOG IgG1 clone 8-18C5 was used as a positive control. Antibodies were IgG1, except in D where three C1 isotypes were tested. **(A)** ELISA against bMOG (left) or wild-type mouse and human MOG_1-125_ (mMOG and hMOG; right). Quantity of bound IgG is expressed as either raw absorbance minus background (left) or absorbance normalized to 8-18C5 (right). **(B)** Western blot for bMOG, mouse MOG_1-125_ or full-length MOG from mouse spinal cord extracts. **(C)** Spinal cord sections stained with anti-MOG antibodies (red) and DAPI (blue). Scale bar: 25 µm. CC = central canal. **(D)** Flow cytometry of live GL261 cells transfected to produce full-length MOG (red), incubated with anti-MOG antibodies, and stained with anti-mouse IgG antibody. Non-transfected cells (blue) were used as a negative control. **(E)** Severity of EAE in mice immunized with MOG_35-55_ and injected intravenously 8 days later with PBS or 200 µg of the indicated antibody. Data from two independent experiments are expressed as either daily scores from the day of disease onset (left) or area under the curve (right). Left panel: *significantly different from PBS (two-way ANOVA with repeated measures using rank-transformed data, *P* < 0.0001; post hoc Mann-Whitney tests, *P* ≤ 0.049). Right panel: *significantly different from the other groups (one-way ANOVA, *P* < 0.0001; post hoc Tukey’s test, *P* ≤ 0.0012). Sample size: 9 PBS; 20 clonotype 1; 20 8-18C5. **(F)** Disease incidence of experiments in E. Log-rank test, *P* = 0.0003. Sample size: 10 PBS; 20 clonotype 1; 20 8-18C5.

To directly demonstrate that C1 is non-pathogenic, we administered this antibody to mice on day 8 post-immunization with MOG_35-55_, i.e. just before the onset of disease and opening of the blood-brain barrier^66,67^. We chose to use MOG_35-55_ instead of bMOG because the former does not induce the production of anti-MOG antibodies (data not shown), which could have had a confounding effect in this experiment. As a positive control, we used the anti-MOG IgG1 clone 8-18C5, which has been repeatedly shown to be pathogenic^55,68–78^. As expected, only 8-18C5 increased EAE severity (**Figure 4E**) and incidence (**Figure 4F**) as compared to the control group. We can therefore conclude that an anti-MOG antibody that is negative in live cell-based assay is not pathogenic, at least by a classic mechanism involving its binding to CNS myelin.

### Pathogenic anti-MOG IgG antibodies are secreted in B cell-dependent EAE

As we could only characterize a few clonotypes out of hundreds, we wondered whether bMOG could nevertheless induce the production of pathogenic anti-MOG antibodies. To answer this question, we first quantified serum anti-MOG antibodies over weeks after bMOG immunization using an isotype-specific ELISA. High titers of anti-MOG IgG1, IgG2b, and IgG2c were detected from day 14 onwards, with IgG1 being clearly the most abundant (**Figure 5A**), corroborating our scRNAseq results (**Figure 3D**). Further analysis of sera on day 14, using an isotype-specific live cell-based assay, revealed the presence of IgG1, IgG2b, and IgG2c antibodies capable of binding to membrane-bound MOG (**Figure 5B**), suggesting that they are pathogenic^11,18,79^. These observations were replicated with sera from mice immunized against hMOG, except that the IgG1/IgG2b ratio was lower (**Figure 5A** and **5B**).

**Figure 5.**
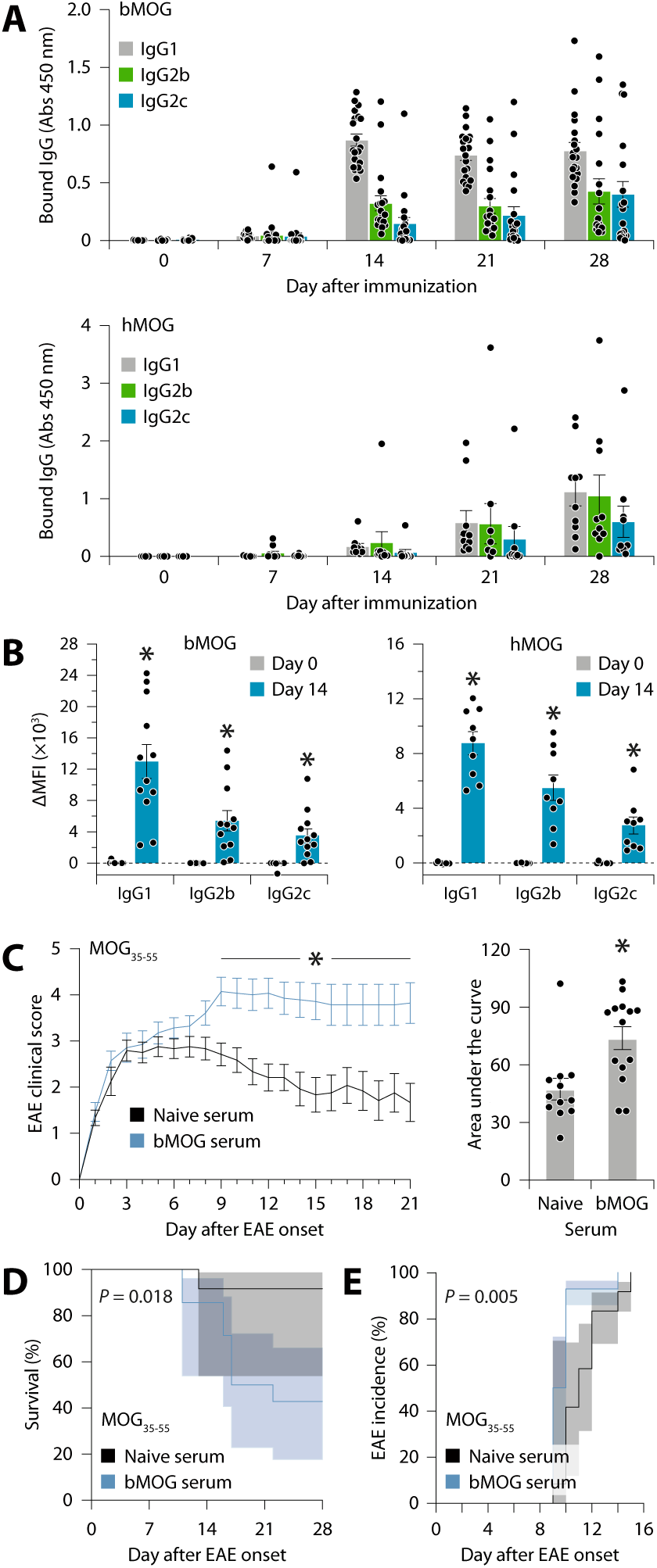
Pathogenic anti-MOG IgG antibodies are produced in B cell-dependent EAE models. **(A)** ELISA detection of anti-MOG IgG1, IgG2b, and IgG2c in serum from mice immunized with either bMOG (top) or hMOG (bottom) through time. Sample size per group: bMOG, 16-20 mice; hMOG, 10 mice. **(B)** Detection of anti-MOG antibodies by live cell-based assay in serum from mice immunized with either bMOG (left) or hMOG (right). GL261 cells transfected to produce full-length mMOG were incubated with serum collected before or after immunization (days 0 and 14), and stained with isotype-specific secondary antibodies. Data are expressed as delta mean fluorescence intensity (ΔMFI). *Significantly different from day 0 group from same isotype (Mann-Whitney test, *P* < 0.0001). Sample size per group: bMOG, 12 mice; hMOG, 9 mice. **(C)** Severity of EAE in MOG_35-55_-immunized mice injected with 150 µl of serum from either naive mice or mice sacrificed 8 days after immunization with bMOG. Data are expressed as either daily scores from the day of disease onset (left) or area under the curve (right). Left panel: *significantly different from naive group as determined by two-way ANOVA with repeated measures using rank-transformed data (*P* = 0.002), followed by post hoc Mann-Whitney tests (*P* ≤ 0.007). Right panel: Mann-Whitney test, *P* = 0.006. Data are from two independent experiments. Sample size: 12 naive serum; 14 bMOG serum. **(D)** Survival and **(E)** disease incidence rates for the experiment in C. *P*-values shown were calculated using the log-rank test. Sample size: as in C.

We next sought to confirm the pathogenicity of bMOG antiserum by adoptive transfer into mice on day 8 post-immunization with MOG_35-55_. As expected, EAE severity was increased in mice administered with bMOG antiserum (**Figure 5C−E**), which is consistent with previous results obtained with hMOG antiserum^55^. Overall, these results indicate that anti-MOG IgG1, IgG2b, and IgG2c are released into the bloodstream from the second week after bMOG immunization and contribute to disease progression.

### 8-18C5 can block the binding of pathogenic antibodies from EAE mice or MOGAD patients

To characterize the binding site of pathogenic bMOG-induced autoantibodies, we performed a competitive live cell-based assay, pre-incubating MOG-expressing cells with 8-18C5 before adding bMOG antiserum. We found that 8-18C5 blocked the binding of anti-MOG IgG2b and IgG2c from all serum samples in a dose-dependent manner (**Figure 6A**). Interestingly, similar results were obtained with serum from hMOG-immunized mice (**Figure 6B**) as well as patients diagnosed with MOGAD and confirmed to be positive for anti-MOG IgG (**Figure 6C**). These findings indicate that pathogenic anti-MOG antibodies, in both mice and humans, bind to MOG by occupying a shared space, and that this binding can be prevented by obstructing this space with a synthetic antibody.

**Figure 6.**
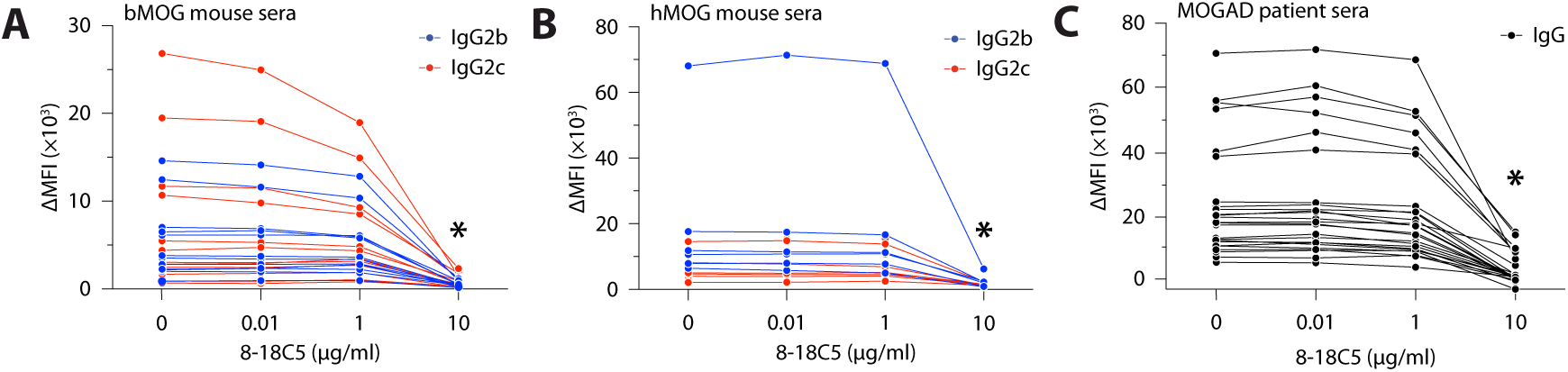
8-18C5 competes with mouse and human pathogenic anti-MOG antibodies for binding to plasma membrane-bound MOG. **(A)** Live cell-based assay in which mMOG-expressing cells were sequentially incubated with 8-18C5 at the indicated concentrations, mouse serum collected on day 14 post-immunization with bMOG, and secondary antibodies to mouse IgG2b and IgG2c. Data are expressed as delta mean fluorescence intensity (ΔMFI). *Significantly different from the other concentrations (Kruskal-Wallis test, *P* ≤ 0.0003 for both IgG2b and IgG2c; post hoc Dunn’s test, *P* ≤ 0.0028). Sample size: 12 sera. **(B)** Same analysis as in A, except that the sera were from hMOG-immunized mice. *Significantly different from the other concentrations (Kruskal-Wallis test, *P* ≤ 0.0072; post hoc Dunn’s test, *P* ≤ 0.0042). Sample size: 6 sera. **(C)** Same analysis as in A, except that hMOG-expressing cells were incubated with serum from MOGAD patients and stained with an anti-human IgG secondary antibody. *Significantly different from the other concentrations (Kruskal-Wallis test, *P* < 0.0001; post hoc Dunn’s test, *P* < 0.0001). Sample size: 25.

### An inactive 8-18C5 variant can attenuate bMOG-induced EAE

Having demonstrated that 8-18C5 competes with pathogenic anti-MOG antibodies, we wondered whether an effector function-deficient 8-18C5 variant could be therapeutically beneficial. To test this, we engineered 8-18C5Mut with the NNAS mutation, which relocates the Fc glycosylation site and blocks binding to FcγRs^80^. This IgG1 variant was unable to bind to mouse or human FcγRs in vitro, unlike 8-18C5, which bound to two mouse and two human FcγRs (mFcγRIIb, mFcγRIII, hFcγRIIa, hFcγRIIb/c), and unlike an irrelevant mouse IgG2a, which bound to all tested FcγRs (**Table 1** and **Supplementary Figure 1**). These results confirm the effect of the NNAS mutation in a mouse antibody context and are consistent with the literature^80,81^.

**Table 1:** Antibody binding affinity to FcγRs determined by surface plasmon resonance.

|  | K<sub>D</sub> (M) |  |  |
| --- | --- | --- | --- |
|  | 8-18C5 (mIgG1) | 8-18C5Mut (mIgG1) | Irrelevant mIgG2 |
| mFcγRI | n/d | n/d | $2.7 \times 10^{-8}$ |
| mFcγRIIb | $3.5 \times 10^{-7}$ | n/d | $6.2 \times 10^{-7}$ |
| mFcγRIII | $2.9 \times 10^{-6}$ | n/d | $3.0 \times 10^{-6}$ |
| mFcγRIV | n/d | n/d | $9.0 \times 10^{-8}$ |
| hFcγRI | n/d | n/d | $6.5 \times 10^{-9}$ |
| hFcγRIIa (R167) | $8.7 \times 10^{-8}$ | n/d | $8.2 \times 10^{-7}$ |
| hFcγRIIb/c | $2.5 \times 10^{-6}$ | n/d | $1.9 \times 10^{-6}$ |
| hFcγRIIIa (V158) | n/d | n/d | $1.7 \times 10^{-6}$ |
| hFcγRIIIa (F158) | n/d | n/d | $2.7 \times 10^{-6}$ |

Next, we administered mice with either 8-18C5Mut, 8-18C5, isotype control IgG1 or PBS on day 9 post-immunization with bMOG. This time point was chosen because it coincides with the increase in blood-brain barrier permeability^66,67,82^ and the appearance of autoantibodies in the bloodstream, which is observable in a fraction of mice as early as day 7 (**Figure 5A**). Interestingly, 8-18C5Mut markedly reduced EAE severity (**Figure 7A**), resulting in a higher rate of complete recovery of locomotor function (**Figure 7B**). Histological examination of spinal cords on day 43 post-immunization revealed clear signs of demyelination (e.g. weak or granular luxol fast blue staining, presence of inflammatory lesions) in mice with a score ≥ 0.5, but not in fully recovered mice (**Figure 7C**). A protective effect of 8-18C5Mut was also observed in another experiment comparing it with C1 (**Supplementary Figure 2**). These results provide proof of concept that a pathogenic autoantibody, once inactivated, can counter the progression of an autoimmune disease by acting as an antagonist.

**Figure 7.**
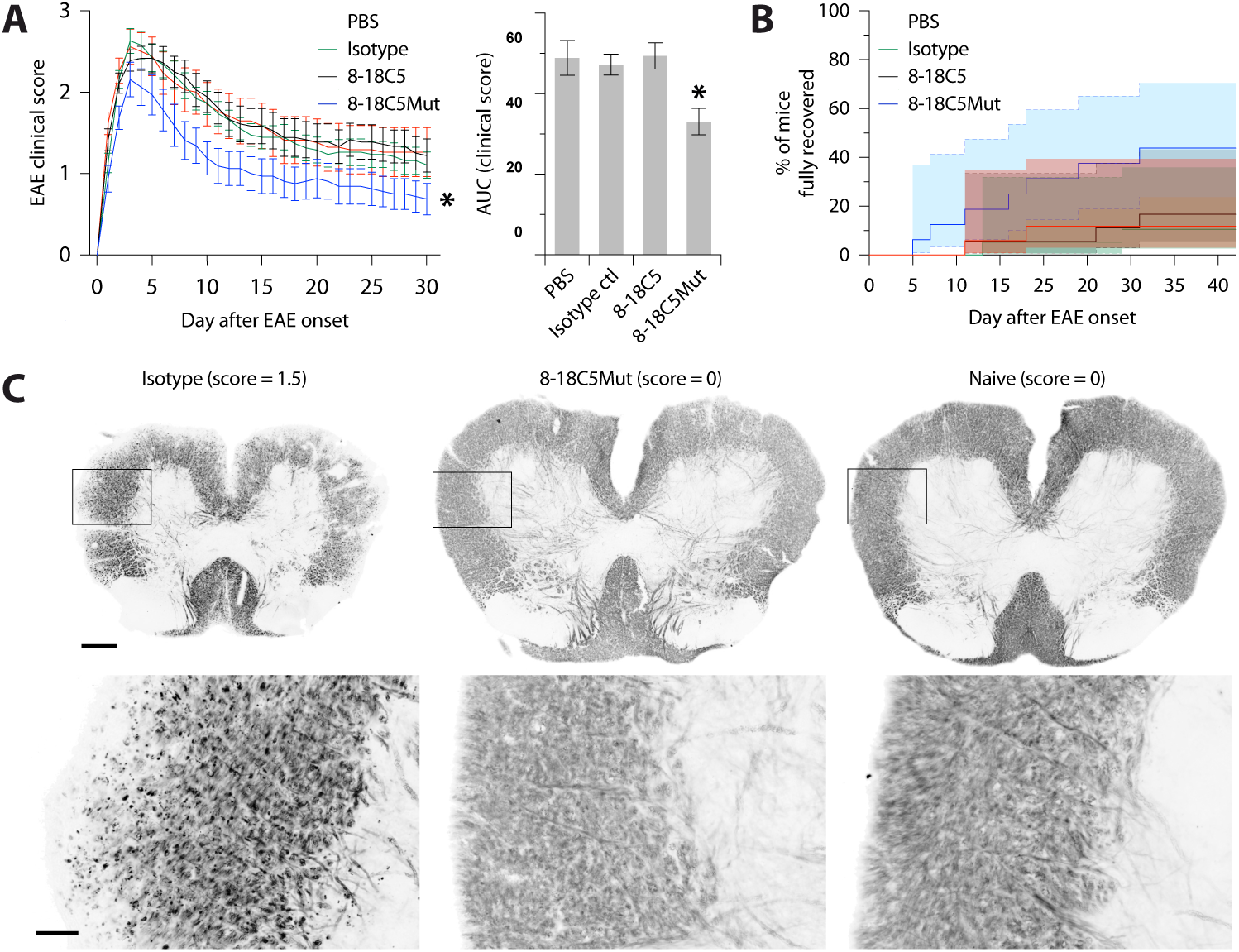
bMOG-induced EAE can be attenuated with 8-18C5Mut. **(A)** Severity of EAE in mice intravenously injected with PBS or 200 µg of the indicated antibody on day 9 post-immunization. Data are expressed as either daily scores from the day of disease onset (left) or area under the curve (right). Left panel: *significantly different from the isotype group, as determined by two-way ANOVA with repeated measures (*P* = 0.049), followed by Fisher’s LSD tests (*P* ≤ 0.0451), using rank-transformed scores. Right panel: *significantly different from the other groups (one-way ANOVA, *P* = 0.0036; post hoc Tukey tests, *P* ≤ 0.0220. Data are from two independent experiments. Sample Size: 18 8-18C5, 16 8-18C5Mut, 19 isotype, 17 PBS. **(B)** Kaplan-Meier plot showing the percentage of mice from panel A that had completely recovered by the end of experiment (log-rank test: overall, *P* = 0.0331; PBS vs 8-18C5Mut, *P* = 0.0412; isotype vs 8-18C5Mut, *P* = 0.0229; 8-18C5 vs 8-18C5Mut, *P* = 0.0680). Shaded areas: 95 % pointwise confidence interval. **(C)** Black gold-stained spinal cord sections on day 43 post-immunization at low (top) or high (bottom) magnification. Evidence of demyelination (lost or granular staining) is observable in a mouse treated with isotype antibody, but not in a mouse treated with 8-18C5Mut or not immunized (naive). The clinical score at the time of sacrifice is indicated for each mouse. Scale bars: top, 200 µm; bottom, 50 µm.

## Discussion

This study aimed to improve our understanding of the humoral response in EAE with a view to finding a strategy to combat autoantibodies. Our results indicate that pathogenic anti-MOG antibodies in B cell-dependent EAE models resemble those in MOGAD patients, targeting a common binding space on the antigen surface. Based on this insight, we came up with the idea of exploiting the NNAS mutation, which is known to inactivate all effector functions of human IgG antibodies^80^. For the first time, we demonstrate the validity of this mutation in a mouse antibody context and, more importantly, its utility in the design of an autoantibody antagonist capable of mitigating an autoimmune disease. More specifically, our inert 8-18C5Mut antibody is directly translatable for the treatment of MOGAD.

The humoral response we observed in bMOG EAE is typical of a primary acute extrafollicular response, in which short-lived plasmablasts massively expand without entering follicular germinal centers and without undergoing affinity maturation, to quickly produce as many antibodies as possible in the short time that they are allotted^2,3,83^. This response bypasses tolerance checkpoints that control the germinal center response^6,84^, thus running a greater risk of generating autoantibodies. This is the first report of a pathogenic extrafollicular response in B cell-dependent EAE. Extrafollicular plasmablasts have been reported in MOG_35-55_ EAE, as we observed here in small numbers, but these cells play an anti-inflammatory role in this B cell-independent model, apparently via IL-10^41,42,85^. Although bMOG-induced plasmablasts also express IL-10, they should not be considered protective as they secrete pathogenic autoantibodies. Rather than acting as a general suppressor of inflammation, IL-10 may contribute to the development of the extrafollicular response, as reported in lupus erythematosus^86^. Furthermore, a pathogenic extrafollicular response is likely to occur in human autoimmune neurological diseases^87^. It may even be favored at the expense of a germinal center response by pathogens such as Epstein-Barr virus^88–91^, which has been linked to autoantibodies in MS^92^. Of course, proving this in humans remains challenging due to the difficulty of obtaining the necessary tissues.

The fate of extrafollicular plasmablasts in our model is unclear. First, some may undergo affinity maturation before differentiating into long-lived plasma cells that would help perpetuate the disease. The increased number of CD138^+^ cells in the bone marrow on day 24 after bMOG immunization supports this possibility. This most likely occurred in two studies that found anti-MOG antibodies with somatic hypermutation in mice immunized against hMOG^55^ or rat cerebellar glycoproteins^68^. In both cases, the mice were immunized several times and sacrificed weeks later. This probably also occurs in people with MOGAD or MS, in whom affinity-matured antibodies have been detected in cerebrospinal fluid^93^. Second, other cells may migrate to meningeal lymphoid follicles, where they would expand to release antibodies inside the CNS. The clusters of CD138^+^ cells that we observed in the leptomeninges 28 days post-immunization may correspond to those reported in ∼40 % of people with secondary progressive MS, which correlate with gray matter demyelination and disease progression^61^.

In conclusion, this study provides the most comprehensive characterization to date of the humoral response occurring in B cell-dependent EAE, which is a more representative model of neurological autoimmune diseases, especially MOGAD, compared with B cell-independent EAE. Our model proves advantageous for studying the mechanisms involved in humoral autoimmunity (e.g. plasmablast development, isotype class switching, somatic hypermutation, ectopic follicle formation, antibody effector functions) as well as for exploring novel therapies targeting them. The NNAS mutation holds promise for the development of autoantibody antagonists, with potential applications extending beyond demyelinating diseases to encompass a range of conditions involving tissue-destructive autoantibodies.

## Materials and Methods

### Human samples

Serum was collected from MOGAD patients (3 male and 2 female children aged 5−17 years, and 10 male and 12 female adults aged 21−72 years) at the Mayo Clinic (Rochester, USA), Hôpital Neurologique Pierre Wertheimer (Bron, France), and University of Montreal Hospital Center (Montreal, Canada) with the approval of the respective ethics committees. All patients gave their informed consent and met the 2023 diagnosis criteria for MOGAD, including a clear positive MOG-IgG cell-based assay on serum^13^.

### Mice

C57BL/6J, Vert-X^56^, CD19-Cre^57^, and Ai14^58^ mice were obtained from The Jackson Laboratory. CD19-Cre and Ai14 mice were crossed to produce CD19^cre^ × Ai14 mice. Genotypes were confirmed by PCR as recommended by the supplier. Experiments were performed on males aged 8 to 12 weeks under specific pathogen-free conditions with the approval of the Laval University Animal Protection Committee and in accordance with the guidelines of the Canadian Council on Animal Care. Groups were formed so that there were no significant differences in age between them.

### EAE induction

Mice received a total of 200 µl of emulsion, injected subcutaneously into each flank. Emulsion was made by mixing equal volumes of Freund’s adjuvant (BD Difco) supplemented with 5 mg/ml of killed *Mycobacterium tuberculosis* H37Ra (BD Difco), and PBS containing 3.0−4.5 mg/ml of either MOG_35-55_ (Medicinal Chemistry Platform, *CHU de Québec*), bMOG, or hMOG. The latter two antigens were produced as previously described^94^. Mice were injected intraperitoneally with 20 µg/kg of pertussis toxin (List Biological Laboratories) immediately before immunization and 2 days later.

### EAE scoring

Mice were weighed and scored daily and blindly as follows: 0, no visual sign of disease; 0.5, partial tail paralysis; 1, complete tail paralysis; 1.5, weakness in one hind limb; 2, weakness in both hind limbs; 2.5, partial hind limb paralysis; 3, complete hind limb paralysis; 3.5, partial forelimb paralysis; 4, complete forelimb paralysis; 5, dead or sacrificed for ethical reasons.

### Administration of serum or antibodies

Serum was collected via cardiac puncture and administered via the retro-orbital sinus at a dose of 150 µl per mouse. Recombinant antibodies (see below) were administered via the retro-orbital sinus at a dose of 200 µg per mouse.

### Recombinant antibody production

Antibodies were cloned and produced in HEK293 cells by MediMab (Montreal) or in CHO cells by Evitria (Zurich) using the variable domain sequences in **Supplementary Table 1**. HEK293 productions were used for in vitro assays, whereas CHO productions were used for in vivo experiments.

### Single-cell RNA sequencing

Single-cell suspensions were prepared from inguinal lymph nodes on day 8 post-immunization with bMOG. CD138^+^ cells were enriched using the EasySep Mouse CD138 Positive Selection Kit (Stemcell Technologies). After determining cell purity (41 % ± 7 SD) and viability (90 % ± 3 SD) by flow cytometry, cells were counted using a TC10 Automated Cell Counter (Bio-RAD) and processed with the Chromium Single Cell Chip A and Chromium Controller (10× Genomics) with the goal of analyzing ∼6,000 cells per mouse. RNA samples were pooled for reverse transcription and amplification to generate gene expression and V(D)J libraries using the Single Cell 5’ Library Kit and Single Cell V(D)J Enrichment Kit (10× Genomics). Libraries were pooled in equimolar ratio and sequenced using both Illumina Hiseq 2500 PE100 technology (low pass) at the University Hospital Center of Quebec and Illumina NovaSeq 6000 S4 PE150 technology (high pass) at Genome Quebec. The mean number of reads per cell was 48,201 (SD ± 8,842) for the gene expression libraries and 16,448 (SD ± 2197) for the V(D)J libraries.

For gene expression libraries, sequencing data were processed, aligned to the mm10 reference genome, and aggregated into a single file using Cell Ranger 4.0 (10× Genomics) with default settings. This file was examined using Loupe Browser 4.2 (10× Genomics) and filtered to remove ambiguous cells (e.g. doublets, phagocytosed cells) based on colocalized cell-specific markers (e.g. *Sdc1* [CD138]*, Ighd, Cd19, Cd3e, C1qa, Siglech, Ccr7, Ly6g, Hbb-bt*). Downstream analyses were performed on the filtered data reanalyzed with Cell Ranger. Cell subsets were identified using tSNE distribution, K-means, and cell type-specific markers. Differentially expressed mRNAs were identified by significant feature comparison. Heatmaps were generated with Clustvis^95^.

For V(D)J libraries, sequencing data were processed and aligned to the GRCm38 reference genome using Cell Ranger with default settings. Each sample was separately examined using Loupe V(D)J Browser v3.0 (10× Genomics). Clonotype and lineage data were obtained using Enclone v.0.5.9 (10× Genomics), first using each sample individually, and then by combining the samples using the *MIX_DONORS* argument to identify interindividual differences and similarities. *MIN_CELLS* was used to keep only clonotypes with ≥ 5 cells, while *PLOT_BY_ISOTYPE* was used to generate honeycomb graphs.

### Flow cytometry

Mice were anesthetized and exsanguinated by transcardial perfusion with saline. Inguinal lymph nodes and spleens were minced, while bone marrow was flushed out from femurs with 10 ml of HBSS (Wisent Bioproducts). The resulting suspensions were filtered through 70-µm cell strainers and treated with RBC lysis buffer (eBioscience). Spinal cords were minced in DPBS containing calcium and magnesium, then digested 45 min at 37 °C in DPBS supplemented with 0.13 U/ml Liberase TM (Roche Diagnostics) and 50 U/ml DNase (Millipore Sigma), filtered through 70-µm cell strainers, and centrifuged for 30 min at room temperature on a 35 % Percoll gradient (GE Healthcare) to remove myelin debris. For immunostaining, cells were incubated on ice for 5 min with rat anti-CD16/CD32 antibody (BD Biosciences, clone 2.4G2, 5 µg/ml) and Fixable Viability Dye eFluor 660 or UV455 (eBioscience, 1:1000), followed by a 30-min incubation with primary antibodies (**Supplementary Table 2**). Cells were washed and resuspended in PBS supplemented with BSA (Cytivia) and EDTA before being analyzed with an LSR II or FACSCanto II flow cytometer (BD Biosciences). For data analysis, the following quality control checks were performed using FlowJo (Tree Star): 1) debris were removed using FSC-A and SSC-A; 2) doublets were removed using FSC-A and FSC-H; and 3) dead cells positive for the viability dye were removed. Gates were based on fluorescence-minus-one controls. Cell counts were normalized to sample volume using 123count eBeads (Thermo Fisher).

### Mass cytometry

Inguinal lymph node cells were isolated as for flow cytometry. For each mouse, 3 × 10^6^ cells were stained with Cell-ID Cisplatin and metal-conjugated antibodies (**Supplementary Table 2**) following the MaxPar Cell Surface Staining with Fresh Fix protocol (Fluidigm). Cells were counted and resuspended at a concentration of 1 × 10^6^ cells/ml in a 9:1 ratio of MaxPar Cell Acquisition Buffer and EQ Four Element Calibration Beads (Fluidigm). Data were acquired using a Helios mass cytometer equipped with a wide-bore injector (Fluidigm), then analyzed using FlowJo (BD Bioscience). Data cleaning was performed according to the technical note Approach to Bivariate Analysis of Data (Fluidigm). After excluding unwanted cells (CD45^−^ cells, TER119^+^ erythrocytes, CD3e^+^ T cells, Ly6G^+^ neutrophils, NK1.1^+^ natural killer cells), data from each mouse were combined to create a single tSNE plot using the following parameters: CD19^+^CD138^−^ B cells limited to 10,000; perplexity of 110; 3,000 iterations; learning rate of 1,889; exact k-nearest neighbors algorithm, Barnes-Hut gradient algorithm.

### ELISA

The binding of recombinant anti-MOG antibodies to MOG antigens was analyzed using in-house ELISAs. Microplates (Corning #9018) were coated overnight at 4 °C with 5 µg/ml of either bMOG (produced in-house) or MOG_1-125_ (Anaspect). BSA-6His-coated wells were used to control for binding to bMOG and MOG_1-125_ tags. The plates were then washed and blocked with 1 % BSA in PBS before adding serial dilutions of recombinant antibodies (50−5000 ng/ml) and incubating for 2 h at room temperature. After washes, a 1:5000 dilution of alkaline phosphatase-conjugated anti-mouse antibody (Jackson Immunoresearch) was added to the plates and incubated 1 h at room temperature. Finally, after additional washes, 1 mg/ml pNPP substrate (MilliporeSigma) was added for up to 30 min at room temperature and absorbance was read at 405 nm using a SpectraMax i3x microplate reader (Molecular Devices). Data shown are those of the weakest dilution for which no signal saturation was observed.

Anti-MOG antibodies were quantified in mouse sera collected via the submandibular vein at a 1:1000 dilution using an anti-mouse MOG_1-125_ ELISA kit (Anaspec). For isotype-specific anti-MOG ELISA, the HRP-conjugated secondary antibody from this kit was replaced by HRP-conjugated goat anti-mouse IgG1, IgG2b, or IgG2c secondary antibody (Abcam). Data are presented as raw absorbance value minus background.

### Western blotting

Spinal cords were collected from saline-perfused mice and homogenized in RIPA buffer (50 mM Tris-HCl, 150 mM NaCl, 1 % Triton X-100, 0.5 % sodium deoxycholate, 0.1 % sodium dodecyl sulfate, 1× protease and phosphatase inhibitor cocktails [Sigma-Aldrich]). Tissue extracts and recombinant MOG samples were diluted, respectively, at 2 mg/ml or 2 ng/ml in Laemmli buffer and boiled for 5 min. Ten µl of each sample were resolved on 10 % SDS-polyacrylamide gel (Bio-Rad Mini-Protean II) and transferred to a nitrocellulose membrane for 1 h at 4 °C and 100 V in transfer buffer (25 mM Tris, 200 mM glycine, 20 % methanol). The membranes were blocked for 30 min at room temperature and incubated overnight at 4 °C in primary antibody, followed by washes and 1 h of incubation at room temperature in HRP-conjugated goat anti-mouse IgG heavy chain (1:5000, ABClonal). Additional washes were done before incubating the membranes 60 sec in Clarity Max ECL substrate (Bio-Rad) and capturing chemiluminescent signal via ChemiDoc XRS+ Imaging System (Bio-Rad).

### Live cell-based assay

The binding of antibodies to membrane-bound MOG was tested using mouse GL261 cells stably transfected with a pCMV vector expressing either full-length mMOG or hMOG isoform 1. Cells were incubated 15 min on ice in PBS supplemented with 2 % FBS and Fixable Viability Dye (eBioscience, 1:1000). For competition assays only, 8-18C5 was included in this first incubation step at 0.01, 1, or 10 µg/ml. Cells were then incubated with either recombinant anti-MOG antibodies (1 µg/ml), mouse serum (1:40 dilution) collected 2 weeks after bMOG immunization, or MOGAD patient serum (1:40) for 30 min. Finally, cells were washed, incubated 15 min with secondary antibodies (**Supplementary Table 2**), and analyzed with a FACSCanto II flow cytometer (BD Biosciences). Non transfected cells and conditions without recombinant primary antibody or serum were used as controls. Prior to analysis, the following quality control checks were performed using FlowJo (Tree Star): 1) debris were removed using FSC-A and SSC-A; 2) doublets were removed using FSC-A and FSC-H; and 3) dead cells positive for the viability dye were removed. Data are presented as mean fluorescence intensity for all live cells, from which the mean fluorescence intensity of control cells was deducted.

### Analysis of antibody-FcγR binding using surface plasmon resonance

Analysis of antibody binding to mouse and human FcγRs was performed on a Biacore T200 (GE Healthcare) using anti-His or anti-mouse Fab capture. Briefly, His-tagged recombinant human and mouse FcγRs (R&D Systems) were diluted to 2 μg/ml in HBS-EP+ (10 mM HEPES, pH 7.4, 150 mM NaCl, 3 mM EDTA, 0.05 % surfactant P20) and captured to anti-tetra His mIgG1 (Qiagen) amine-coupled to a CM5 chip (Cytiva) for 30 sec at a flow rate of 10 μl/min. Antibodies were serially diluted 2- or 3-fold from 3000 nM and injected over the captured FcγRs for 1 min in duplicate at 30 μl/min. To measure binding to human FcγRI, antibodies were serially diluted 2-fold from 300 nM and injected for 3 min followed by 10 min dissociation. The surface was regenerated with 10 mM glycine, pH 1.5. Alternately, human and mouse FcγRII binding was measured by capturing ∼100 RU of antibody to a CM5 chip immobilized with goat anti-mouse IgG, F(ab’)₂ (Jackson ImmunoResearch) and injecting receptors serially diluted 2-fold from 3000 nM in HBS-EP+ buffer for 1 min at 30 μl/min. Dissociation was measured for 1 min and the surface was regenerated with 0.85 % phosphoric acid. Steady state analysis was used to determine the binding affinity of low affinity receptors and kinetic fits using a 1:1 binding model were performed on high affinity receptor sensorgrams to calculate K_D_.

### Histology

For lymph nodes, mice were transcardially perfused with saline for 5 min. Lymph nodes were fixed overnight at 4 °C in PLP solution (1 % PFA, 0.1M L-lysin, 0.01 M sodium periodate in PBS), and mounted in blocks of M-1 Embedding Matrix (Thermo Fisher). Series of 8 µm-thick sections were cut using a cryostat (CryoStar NX70), mounted on slides, and stored at −80 °C.

For spinal cords, mice were transcardially perfused with ice-cold saline, followed by 4 % PFA in phosphate buffer, pH 7.4, over 10 min. Spinal cords were post-fixed 4 h at 4 °C in 4 % PFA and cryoprotected overnight in 20 % sucrose. 25 µm-thick sections were cut with a microtome (Leica SM2000R) and stored at −20 °C in cryoprotectant solution (PBS with 20 % glycerol and 30 % ethylene glycol).

Immunofluorescence was performed as previously described^96^. Briefly, slides were blocked and incubated overnight at 4 °C in primary antibodies (**Supplementary Table 2**). After washing, sections were incubated at room temperature for 2 h in secondary antibodies (**Supplementary Table 2**). Slides were counterstained for 1 min with 2 µg/ml DAPI, then mounted with coverslips no. 1.5H and ProLong Glass Antifade Mountant (Molecular Probes).

Sections of lumbar spinal cords were stained for myelin with 0.3 % Black-Gold II (Histo-Chem) in 0.9 % NaCl at 60 °C for 15 min, then differentiated in 1 % sodium thiosulphate at 60 °C for 3 min as described^97^. Brightfield tiled images were captured at 20× magnification using a Zeiss Axio Scan.Z1 slide scanner.

### Confocal microscopy

Confocal images were acquired with a Leica TCS SP8 STED 3X microscope by sequential scanning using the following settings: objective, HC/PL/APO 63×/1.40 oil; immersion oil, Leica Type F; scan speed, 600 Hz; line average, 2-4; time gate, 0.3-6.0 ns. Laser power and gain were set to optimize signal-to-noise ratio and avoid saturation using the QLUT Glow mode. Sizes of pixel, pinhole and *z*-step were set to optimize resolution or to oversample in the case of images to be deconvolved. Deconvolution was performed with Huygens Professional (Scientific Volume Imaging) using a theoretical point spread function, manual settings for background intensity and default signal-to-noise ratio. Color balance, contrast and brightness were adjusted with Photoshop (Adobe). CD138^+^ and CD3^+^ cells were systematically counted in the meninges of five serial sections per mouse.

### Statistics

Data are expressed as mean ± standard error. In general, groups were compared using either nonparametric (Mann-Whitney, Kruskal-Wallis with Dunn’s test) or parametric tests (one-way ANOVA, two-way ANOVA with Tukey’s or Dunnett’s test), when data were continuous, normally distributed (Anderson-Darling test), and of equal variance (Levene’s test). EAE incidence curves were constructed using the Kaplan-Meier method and compared by log-rank analysis. EAE severity curves were compared by: 1) two-way ANOVA with repeated measures, followed by Fisher’s LSD test, using rank-transformed scores; and 2) parametric tests using the area under the curve. Analyses were performed with GraphPad Prism 9.1 (GraphPad Software). A *P* value less than 0.05 was considered significant.

## Data availability

The raw and processed scRNAseq data generated in this study are available in the NCBI GEO repository under the accession code <u>GSE260585</u>.

## Acknowledgments

We thank Catherine Bélanger, Aline Dumas, and Alexandre Patenaude for initial experiments that have led to this study. We thank Dr. Jennifer Gommerman (University of Toronto, Canada) for the gift of the hMOG plasmid. We thank Drs. Nathalie Arbour and Alexandre Prat (University of Montreal Hospital Center) as well as Nathalie Dufay, Margaux Millet, and Lakhdar Benyahya (NeuroBioTec, Hospices Civils de Lyon, Lyon, France) for access to samples from the biobanks BH07.001 and BB-0033-00046, respectively. We are grateful to the staff of the *CHU de Québec* Genomic Platform and *Génome Québec* for RNA library preparation and sequencing. We thank the *CHU de Québec* high-throughput image analysis platform for assistance with slide scanning. We thank the staff of Medimab (Montreal) and Evitria (Zurich) for the production of antibodies. This work was supported by grants to L.V. from the Canadian Institutes for Health Research, the Multiple Sclerosis Society of Canada, and the Natural Sciences and Engineering Research Council of Canada. It was also supported by the Canada Research Chair in Medical Genomics and Grant R01NS113828 from the National Institute of Neurological Disorders and Stroke, respectively awarded to J.C. and E.P.F.

## Author contributions

L.V. was responsible for conceptualization, funding acquisition, project administration, supervision, methodology, data analysis, and writing. R.T.M. carried out most of the experiments and contributed to data analysis and writing with the help of F.M. Y.Z. performed the mass cytometry experiment, while L.B. performed the analysis of plasmablasts in spinal cord sections, with the help and supervision of L.C.B. Q.Z, J.J., and A.P. provided the NNAS sequence and performed the antibody-FcγR binding analysis. J.M.D. processed and analyzed scRNAseq data under the supervision of J.C. E.F., R.M., and C.L. provided human serum samples. S.K. contributed to conceptualization and provided bMOG. All authors reviewed the manuscript and gave final approval.

## Supplementary Materials

**Supplementary Dataset 1.** scRNAseq of bMOG-induced plasmablasts.

**Supplementary Figure 1.** Surface plasmon resonance sensorgrams comparing the binding of mouse antibodies to mouse and human FcγRs.

**Supplementary Figure 2.** bMOG-induced EAE can be attenuated with 8-18C5Mut.

**Supplementary Table 1.** Antibody clones selected for further analysis.

**Supplementary Table 2.** Antibodies used in this study.

