## Supplementary Figure 1 for "Turncoat antibodies unmasked in a model of autoimmune demyelination: from biology to therapy"

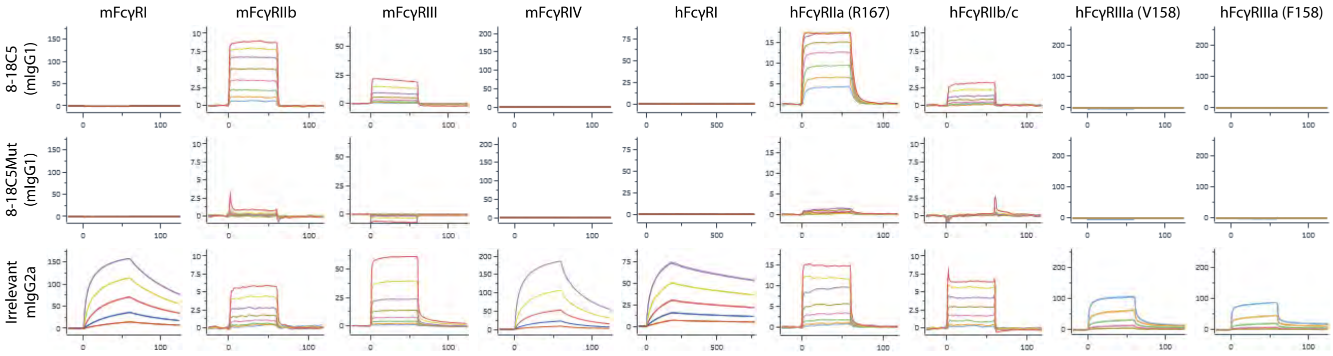


**Supplementary Figure 1.** Surface plasmon resonance sensorgrams comparing the binding of mouse antibodies to mouse and human FcγRs. The relative change in binding response (resonance units) for various concentrations of antibodies is shown over time (seconds). $K_D$ values are shown in Table 1.
