## Supplementary Figure 2 for "Turncoat antibodies unmasked in a model of autoimmune demyelination: from biology to therapy"

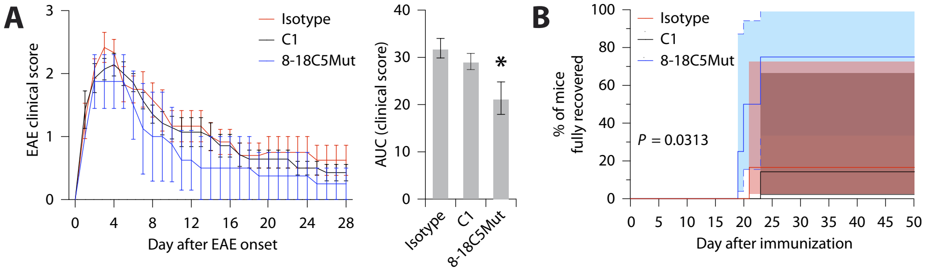


**Supplementary Figure 2.** bMOG-induced EAE can be attenuated with 8-18C5Mut. **(A)** Severity of EAE in mice intravenously injected with 200 µg of the indicated antibody on day 1 post-immunization. Data are expressed as either daily scores from the day of disease onset (left) or area under the curve (right). *Significantly different from the isotype control group (one-way ANOVA, $P$ = 0.0239; post hoc Dunnett's test, $P \leq 0.0149$. **(B)** Kaplan-Meier plot showing the percentage of mice that had completely recovered by the end of experiment (log-rank test: overall, $P$ = 0.0313. Shaded areas: 95 % pointwise confidence interval. Sample size: 6 C1, 4 8-18C5Mut, 6 isotype.
