## Supplementary Table 2 for "Turncoat antibodies unmasked in a model of autoimmune demyelination: from biology to therapy"

**Supplementary Table 2**. Antibodies used in this study.

| Antigen | Clone | Host | Conjugate | Working conc. (µg/ml) | Company |
|---|---|---|---|---|---|
| **Flow Cytometry** | | | | | |
| CD3ε | 145-2C11 | Hamster | PE, PE-Cy5, PE-Cy7 | 2 | BD Biosciences, Biolegend, Thermo Scientific |
| CD4 | RM4-5 | Rat | PerCP-Cy5.5 | 1 | BD Biosciences |
| CD8A | APC | Rat | 53-6.7 | 1 | BD Biosciences |
| CD11b | M1/70 | Rat | eFluor 450, APC-Fire 750 | 1 | ebiosciences, Biolegend |
| CD11c | N418 | Hamster | BV510, PE-Cy5 | 1 | Biolegend |
| CD16/32 | 2.4g2 | Rat | - | 5 | BD Biosciences |
| CD19 | 1D3 | Rat | eFluor 660, PE-Cy7 | 1 | ebiosciences |
| CD45 | 30-F11 | Rat | PerCP-Cy5.5, BV605, BV510 | 1 | Biolegend |
| CD138 | 281-2 | Rat | PE-CF594 | 1 | Biolegend |
| **Mass cytometry** | | | | | |
| CCR7 | 4B12 | Rat | 155Gd | 2 | Biolegend |
| CD11b | M1/70 | Rat | 148Nd | 0.5 | Fluidigm |
| CD11c | N418 | Hamster | 142Nd | 2 | Fluidigm |
| CD3ε | 145-2C11 | Hamster | 152Sm | 1 | Fluidigm |
| CD19 | 6D5 | Rat | 149Sm | 0.25 | Fluidigm |
| CD45 | 30-F11 | Rat | 89Y | 0.5 | Fluidigm |
| CD70 | AP-MAB0825 | Rat | 158Gd | 0.5 | Novus Biologicals |
| CD80 | 16-10A1 | Hamster | 171Yb | 2 | Fluidigm |
| CD83 | Polyclonal | Goat | 153Eu | 4 | R&D Systems |
| CD86 | GL1 | Rat | 172Yb | 1 | Fluidigm |
| CD138 | 281-2 | Rat | 147Sm | 4 | Biolegend |
| CD209A | MMD3 | Mouse | 156Gd | 4 | Biolegend |
| CSF1R | AFS98 | Rat | 144Nd | 1 | Fluidigm |
| CX3CR1 | SA011F11 | Mouse | 164Dy | 1 | Fluidigm |
| FCER1A | Mar-1 | Hamster | 176Yb | 0.5 | Fluidigm |
| ICAM1 | YN1/1.7.4 | Rat | 163Dy | 1 | Fluidigm |
| KLRA2 | Polyclonal | Rabbit | 175Lu | 4 | Novus Biologicals |
| Ly6G | 1A8 | Rat | 141Pr | 0.5 | Fluidigm |
| MARCO | 2359A | Rabbit | 170Er | 0.5 | R&D Systems |
| MHCII | M5/114.15.2 | Rat | 209Bi | 0.5 | Fluidigm |
| NGFR | Polyclonal | Goat | 166Er | 4 | R&D Systems |
| NK1.1 | PK136 | Mouse | 165Ho | 1 | Fluidigm |
| SIGLEC-H | 551 | Rat | 151Eu | 0.5 | Biolegend |
| SIRPB1 | 10 | Rabbit | 159Tb | 4 | Invitrogen |
| TER-119 | TER-119 | Rat | 154Sm | 2 | Fluidigm |

| | | | | | |
|---|---|---|---|---|---|
| XCR1 | ZET | Mouse | 174Yb | 4 | Biolegend |
| **Immunofluorescence** | | | | | |
| CD3ε | 145-2C11 | Hamster | - | 2 | Biolegend |
| CD138 | 281-2 | Rat | - | 1 | Biolegend |
| Hamster IgG | Polyclonal | Polyclonal | Alexa 488 | 4 | Jackson Immunoresearch |
| LYVE1 | Polyclonal | Rabbit | - | 5 | Fitzgerald |
| Mouse IgG | Polyclonal | Goat | RRX | 2 | Jackson Immunoresearch |
| Rabbit IgG | Polyclonal | Goat | Alexa 488 | 4 | Thermo Fisher Scientific |
| Rat IgG | Polyclonal | Goat | Alexa 633 | 4 | Invitrogen |
| **Live cell-based assay** | | | | | |
| Mouse IgG | Polyclonal | Goat | Alexa 488 | 4 | Thermo Fisher Scientific |
| Mouse IgG1 | Polyclonal | Goat | PE | 1 | Thermo Fisher Scientific |
| Mouse IgG2b | Polyclonal | Goat | APC-Cy7 | 1.5 | Abcam |
| Mouse IgG2c | Polyclonal | Goat | APC | 1 | SouthernBiotech |
| Human IgG | Polyclonal | Goat | Alexa 647 | 5 | Jackson Immunoresearch |
| **ELISA** | | | | | |
| Mouse IgG1 | Polyclonal | Goat | HRP | 0.2 | Abcam |
| Mouse IgG2b | Polyclonal | Goat | HRP | 0.2 | Abcam |
| Mouse IgG2c | Polyclonal | Goat | HRP | 0.2 | Abcam |
| Human IgG | Polyclonal | Goat | HRP | 0.1 | Jackson Immunoresearch |
